# Neutralization Fingerprinting Technology for Characterizing Polyclonal Antibody Responses to Dengue Vaccines

**DOI:** 10.1101/2021.12.02.467518

**Authors:** Nagarajan Raju, Xiaoyan Zhan, Subash Das, Lovkesh Karwal, Hansi J. Dean, James E. Crowe, Robert H. Carnahan, Ivelin S. Georgiev

## Abstract

Dengue is a major public health threat. There are four serotypes of dengue virus (DENV), therefore efforts are focused on development of safe and effective tetravalent DENV vaccines. While neutralizing antibodies contribute to protective immunity, there are still important gaps in understanding of immune responses elicited by dengue infection and vaccination, including defining immune correlates of protection. To that end, here we present a computational modeling framework for evaluating the specificities of neutralizing antibodies elicited to tetravalent DENV vaccines, based on the concept of antibody-virus neutralization fingerprints. We developed and applied this framework to samples from clinical studies of TAK-003, a tetravalent vaccine candidate currently in phase 3 trials, to characterize the effect of prior dengue infection (baseline) on the specificities of vaccine-elicited antibody responses. Our results suggested a similarity of neutralizing antibody specificities in baseline-seronegative individuals. In contrast, amplification of pre-existing neutralizing antibody specificities was predicted for baseline-seropositive individuals, thus quantifying the role of immunologic imprinting in driving antibody responses to DENV vaccines. The analysis framework proposed here can apply to studies of sequential dengue infections and other tetravalent DENV vaccines and can contribute to understanding dengue immune correlates of protection to help guide further vaccine development and optimization.

## Introduction

Dengue viral infection is a serious global health threat, with an estimated hundreds of millions of cases every year worldwide, caused by four different DENV serotypes [1–3]. Although an initial infection with one serotype can induce protective immunity against subsequent infections of the same serotype, prior infection also can lead to antibody-dependent enhancement (ADE) upon infection with a different serotype, which can increase the likelihood of higher disease severity [4, 5]. Hence, tetravalent vaccine formulations that include representative strains from each serotype are a major target for preclinical and clinical development [6, 7]. While the tetravalent Dengvaxia vaccine is already approved for use in certain geographic regions [6, 8-10], no vaccine has been shown to offer complete protection against all four DENV serotypes [10]. Another vaccine candidate, TAK-003, currently being evaluated in a Phase 3 clinical trial (NCT02747927), comprises an attenuated DENV-2 and three recombinant viruses containing the structural premembrane (prM) and the viral envelope (E) proteins of DENV-1, −3, and −4 cloned into the attenuated DENV-2 backbone [11]. TAK-003 demonstrated efficacy against symptomatic dengue in both baseline seronegative and seropositive populations for 3 years postvaccination [11, 12]. TAK-003 elicits type-specific and cross-reactive antibodies in baseline seronegative vaccine recipients [13, 14]. However, little is currently understood about how immune responses to prior infection may affect (imprint on) the profile of antibodies elicited to subsequent vaccination. Neutralizing antibody titers are generally higher in baseline-seropositive individuals, who experienced DENV infection prior to TAK-003 vaccination, compared to baseline-seronegative individuals, who do not have DENV-specific neutralizing antibodies at the time of vaccination [11]. However, studying the specificity of neutralizing antibodies after vaccination of baseline-seropositive individuals is complex. These results highlight the ability of prior infection to serve as an immunologically priming stimulus for DENV-specific antibody responses to a multivalent vaccine.

The characterization of the collective specificities that comprise polyclonal antibodies is a complex process, due to the large diversity of antibody epitopes and phenotypes in response to infection or vaccination for a given pathogen. To that end, previously we had developed a computationally driven approach, designated neutralization fingerprinting (NFP), for deconvoluting polyclonal antibody specificities in the context of HIV-1 infection, through mathematical analysis of antibody-virus neutralization data [15]. In essence, the NFP algorithm compares a polyclonal neutralization pattern over a set of diverse HIV-1 strains to the corresponding patterns for broadly neutralizing reference antibodies, to predict the overall prevalence of each of these reference specificities in the given sample[15–17]. Due to the similarly high antigen sequence diversity for both HIV-1 and DENV, we set out to develop a similar NFP approach for DENV. Application of this approach to clinical samples from TAK-003 suggested that immunologic imprinting plays an important role for defining the types of antibodies elicited to vaccination. Overall, the NFP platform developed here for DENV can be of general utility for deconvoluting the antibody specificities in polyclonal responses to sequential dengue infections and to multivalent vaccines, such as TAK-003.

## Results

### Neutralization fingerprints of DENV-specific monoclonal antibodies

To interrogate the patterns of virus neutralization by monoclonal antibodies, we selected a set of 18 monoclonal antibodies with known antigen specificities and tested their ability to neutralize 25 representative virus strains including strains from all four DENV serotypes (**Figure 1A**). Several diverse antibody neutralization patterns, or fingerprints, were observed, including antibodies that recognized only one of the four DENV serotypes, as well as cross-reactive antibodies that neutralized strains from all four serotypes (**Figure 1A**). In the case of HIV-1, the neutralization fingerprints of antibodies targeting similar epitopes on the virus had high correlations, whereas the correlations for the fingerprints of antibodies targeting different epitopes were generally low [15]. In the case of DENV, however, this pattern of correlations is challenged by the existence of DENV serotypes and serotype specific antibodies (**Figure 1B**). For example, DENV-1-specific antibodies would have the same large number of resistant strains (other than DENV-1), resulting in a high correlation of the DENV-1-specific antibody fingerprints, regardless of any differences in epitopes. Indeed, within our neutralization data, the fingerprints for DENV-1-specific monoclonal antibodies 1F4 and 4L5 exhibited a high correlation (0.92), despite differences in epitope (**Figure 1B**). To address this challenge, in addition to considering epitopes, DENV serotype specificity was included as a criterion for grouping antibodies by neutralization fingerprints, resulting in 7 antibody groups, including the 4 type-specific groups (1F4/4L5-like, 2D22/3F9-like, 5J7-like, 126/131-like) as well as 3 crossreactive groups (C10-like, FL-like, 2M2-like), which describe specific epitopes or domains (**Figure 1C**). Antibody 1L12 appeared to be an outlier and was excluded from further analysis. While it is believed to be a DENV-2-specific antibody [18], the neutralization analysis suggested broad cross-reactivity, with neutralization for non-DENV-2 dengue variants being low-potency and therefore less robust/reliable. Hence, the final dataset contained 17 DENV monoclonal antibodies clustered into 7 antibody groups (**Figure 1C**). As expected, the correlations of the neutralization fingerprints for antibodies within the same group were significantly higher (p<0.0001, t-test) than correlations between antibodies from different groups (**Figure 1D**), confirming the ability to discriminate between the 7 antibody groups based on neutralization fingerprints.

**Figure 1:**
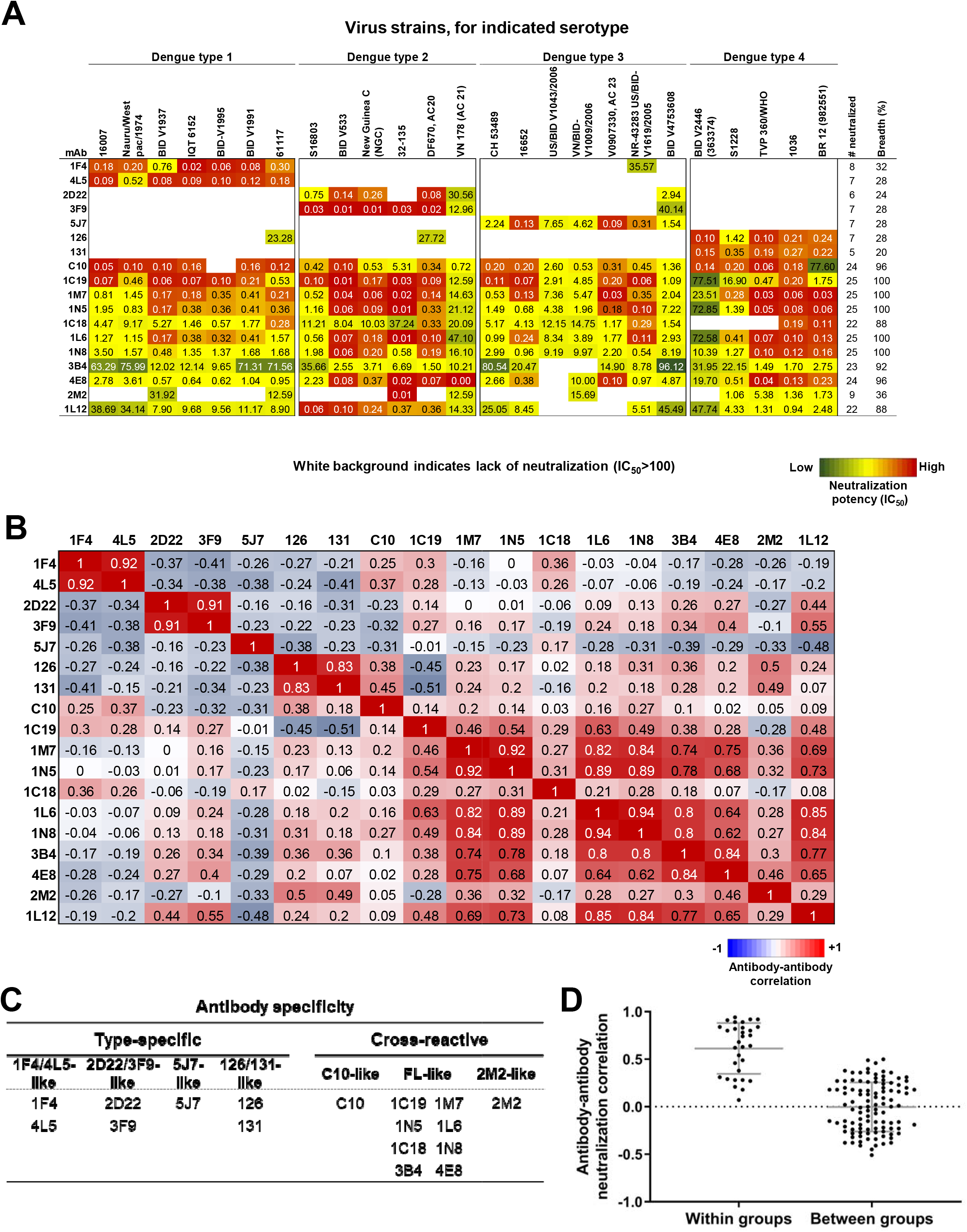
mAb neutralization data and specificity groups. A. Antibody-virus neutralization matrix. Shown are the neutralization IC50 values for 18 antibodies against 25 DENV strains. Color scale is green(low)-yellow-red(high) neutralization; IC50 values greater than 100 are shown in white. Last two columns show the neutralization breadth (number and percent of viruses, respectively) for each antibody. B. Antibody-antibody neutralization correlation matrix. Spearman correlations were computed for the neutralization fingerprints for each pair of antibodies [range: blue (−1) – white (0) – red (+1)]. C. Grouping of DENV antibodies based on two criteria: serotype specificities (1F4/4L5-, 2D22/3F9-, 5J7- or 126/131-like) and cross-reactive specificities (C10-, FL- or 2M2-like). D. Antibody-antibody correlation values within antibody specificity groups (left) and between groups (right).

### Computational selection of an optimized virus panel for polyclonal neutralization analysis

Since the initial 25-virus panel was not selected based on specific metrics other than ensuring the presence of diverse strains from the four DENV serotypes, we next sought to optimize the composition of the virus panel, similarly to our prior work with HIV-1 [17]. The goal of these efforts was to identify subsets of the original 25-virus panel that were associated with improved prediction accuracy for delineating the antibody specificities in polyclonal samples from DENV vaccines studies. To evaluate the prediction accuracy of candidate virus panels, we simulated the polyclonal “serum” neutralization by combinations of pairs of antibodies, each from a different antibody group. To that end, we generated a set of 2,100 simulated polyclonal sera (**Supplementary Figure 1A**) to assess how well the NFP algorithm can predict the known antibody composition in the set of simulated sera, for each candidate virus panel. The prediction accuracy of a given virus panel was measured as the average serum delineation error among all simulated sera, with lower values corresponding to greater accuracy. All possible panels of size 18 to 22 viruses were evaluated, with a number of panels of different size showing improved prediction accuracy compared to the full 25-virus panel (**Figure 2A,B**).

**Figure 2:**
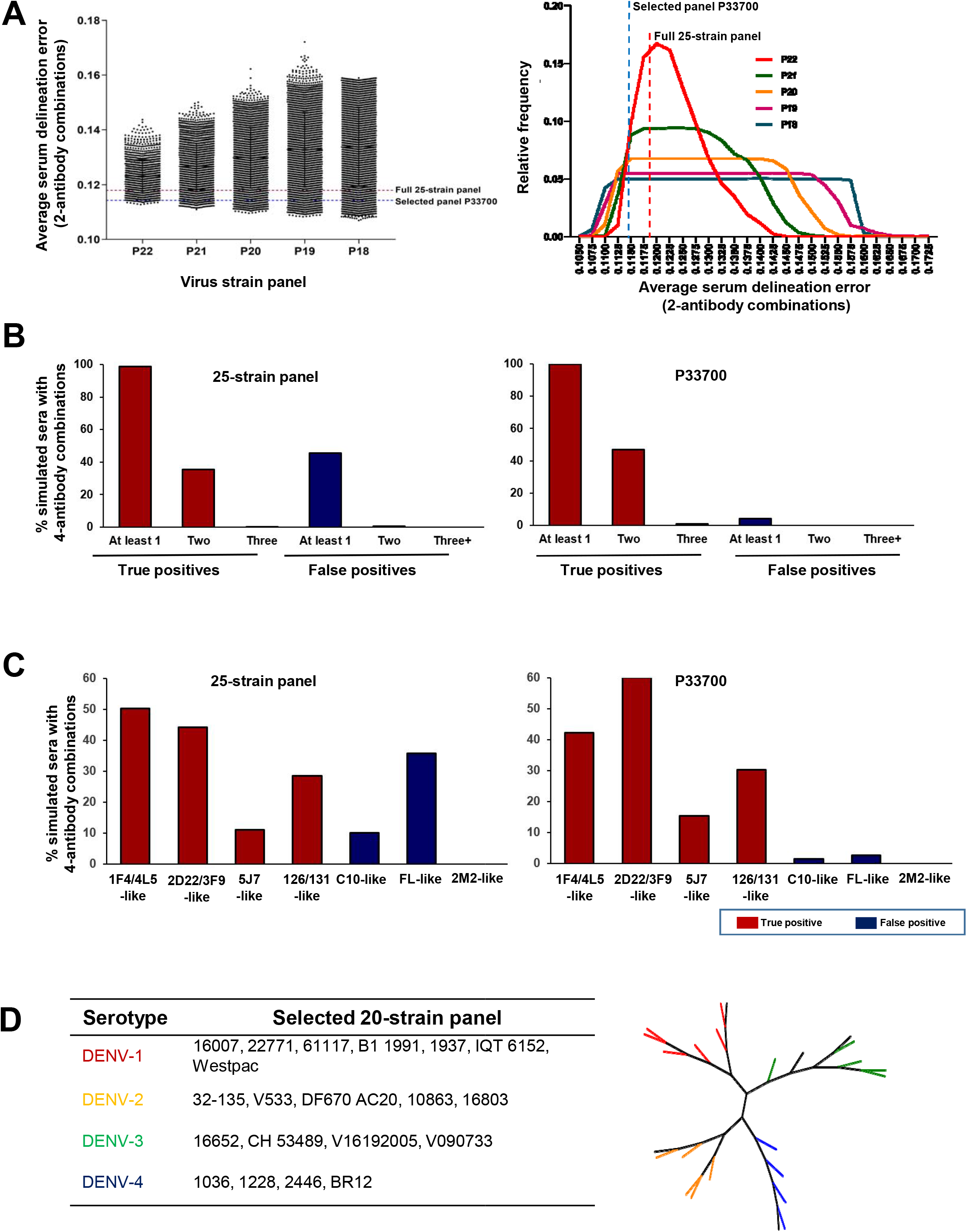
Establishment of neutralization fingerprinting framework - virus panel optimization. A. Scatter plot (left panel) and distribution (right panel) of average serum delineation errors for the simulated sera with 2-antibody combinations for virus panels of different sizes; highlighted are the full 25-strain panel and the selected 20-strain virus panel P33700. B. Bar graphs of true positives (red) and false positives (blue) for the 25-strain (left) and selected 20-strain (right) panel against simulated sera with 4-antibody combinations. C. Bar graphs for the antibody group-wise true positives (red) and false positives (blue) for the 25-strain (left) and selected 20-strain (right) panel against simulated sera with 4-antibody combinations. D. (left) Strain names in the selected panel from each DENV serotype are shown. (right) The selected strains are highlighted with DENV serotype-specific colors in the phylogenetic tree of the full 25-strains.

These virus panels were evaluated further on an independent set of simulated sera that included 4-antibody combinations from the 4 DENV type-specific antibody groups (**Supplementary Figure 1B**). These 4-antibody simulated sera aimed at addressing the question of whether the NFP algorithm can successfully identify polyclonal neutralization breadth as a combination of multiple type-specific but not broadly neutralizing monoclonal antibodies, as opposed to a single broadly neutralizing cross-reactive monoclonal antibody. Based on this analysis, the 20-strain virus panel P33700 was selected for NFP.

### Comparison of NFP prediction accuracy between virus panel P33700 and the full 25-virus panel

Several metrics were visualized to compare the NFP prediction accuracy between the selected 20-virus panel P33700 and the full 25-virus panel (**Figure 2, Supplementary Figure 1**). For the 2-antibody simulated sera, the average serum delineation error was 0.1143 for the 25-strain panel and 0.1180 for the P33700 panel (**Figure 2A**). The two panels had similar true positive rates, with both panels being able to identify at least one of the two specificities for virtually all simulated sera (**Supplementary Figure 1C**). P33700 identified both component antibody specificities in a marginally larger fraction of simulated sera, although both panels could identify two specificities in ~60% of sera (**Supplementary Figure 1C**). A more substantial improvement was observed with the false positive rates, with P33700 resulting in ~50% fewer false positive results than the full 25-virus panel (**Supplementary Figure 1C**). Together, these data suggest that the improved accuracy of P33700 compared to the 25-virus panel on the 2-antibody simulated sera resulted primarily from an improvement in false-positive rates, leading to better serum delineation errors.

Notable differences in the accuracy of the two virus panels were observed for the 4-antibody simulated sera (**Figure 2B**). While in both cases at least one specificity was identified correctly for virtually all simulated sera, the P33700 20-virus panel correctly identified two or more specificities in 48% of simulated sera, an improvement of ~12% compared to the 25-strain panel. Of note, there was a more than 10-fold improvement in the false-positive rates for panel P33700, resulting in a false positive rate of ~4%, compared to ~45% for the full 25-virus panel (**Figure 2B**). To better understand the improvements in prediction accuracy for panel P33700, we compared the true- and false-positive rates for each of the different antibody groups for the 4-antibody simulated sera (**Figure 2C**). While the true-positive rate for 1F4/4L5-like antibodies was lower for P33700, the overall true-positive rate improvement was driven by large improvements for 2D22/3F9- and 5J7-like true-positive rates. The difference in false-positive rates between the two panels could be explained by the substantially decreased false positive results for C10-like and FL-like antibodies (**Figure 2C**). Together, these results suggested that the P33700 panel could more accurately deconvolute polyclonal from monoclonal neutralization breadth and should be better suited for application to DENV antibody response characterization that aims to discriminate between type-specific and cross-reactive antibodies. Based on these data, the 20-virus panel P33700, comprised of 6 strains from dengue serotype 1, 5 from dengue serotype 2, 5 from dengue serotype 3, and 4 from dengue serotype 4 (**Figure 2D**), was selected for evaluating polyclonal antibody responses to the TAK-003 vaccine.

### Neutralizing antibodies elicited to TAK-003 vaccination in baseline negative vs. baseline positive individuals

To evaluate the applicability of the NFP algorithm to human polyclonal sera, we assessed serum neutralization on the P33700 panel for a set of 20 donors vaccinated with TAK-003 (**Figure 3A**). Cohort 1 included samples of 10 baseline-seronegative donors at 30 days postvaccination with TAK-003; we will refer to this group of samples as N30. In contrast, Cohort 2 included 10 baseline-seropositive donors with prior DENV infection, sampled pre-vaccination (P0) and 30 days post-vaccination (P30). To evaluate the differences between the three groups of samples, the geometric mean of the neutralization ID_50_ for all viruses from a given DENV serotype were computed for each serum sample (**Figure 3B**). In the N30 group, neutralization against DENV-2-strains was the strongest, although neutralization against all dengue types was generally observed. This finding is consistent with phase 3 trial results demonstrating that TAK-003 elicits a high rate of tetravalent neutralizing antibodies in donors without prior infection [11]. In the P0 group, neutralization titers were observed for all DENV types (**Figure 3B**). The P0 neutralization titers establish baseline (pre-vaccination) neutralization potencies that can be compared to the neutralization titers post-vaccination in the P30 group. In P30, increased overall neutralization was observed for all DENV types compared to P0, suggesting that TAK-003 boosted neutralizing antibody responses elicited by prior infection with all DENV serotypes (**Figure 3B**). To quantify the neutralization differences between the three sets of samples, statistical significance was evaluated using the geometric mean of ID_50_ values (**Figure 3C**). As P0 and P30 are donor-matched, non-parametric paired correlation analysis was performed, whereas non-parametric non-paired analysis was performed for the comparisons involving N30. When comparing N30 vs. P0, significant differences were not observed for DENV types 1, 3, or 4. However, a significant difference was observed for DENV type 2, indicating a stronger DENV type 2 response to vaccination compared to prior infection. When comparing N30 to P30, significance was observed for neutralization of both DENV-1 and DENV-3, indicating prior infection serves as a strong prime for these DENV serotypes, compared to vaccination of DENV-naïve individuals. Conversely, the differences for DENV-2 and DENV-4 were not significant, indicating that prior infection was not beneficial for developing greater neutralizing antibody titers to vaccination against these DENV serotypes. When comparing the within-donor changes in neutralization potency (P0 vs. P30), significance or trends toward significance were observed for all the DENV serotypes, highlighting the improvement of neutralization in response to vaccination for all DENV serotypes (**Figure 3C**).

**Figure 3:**
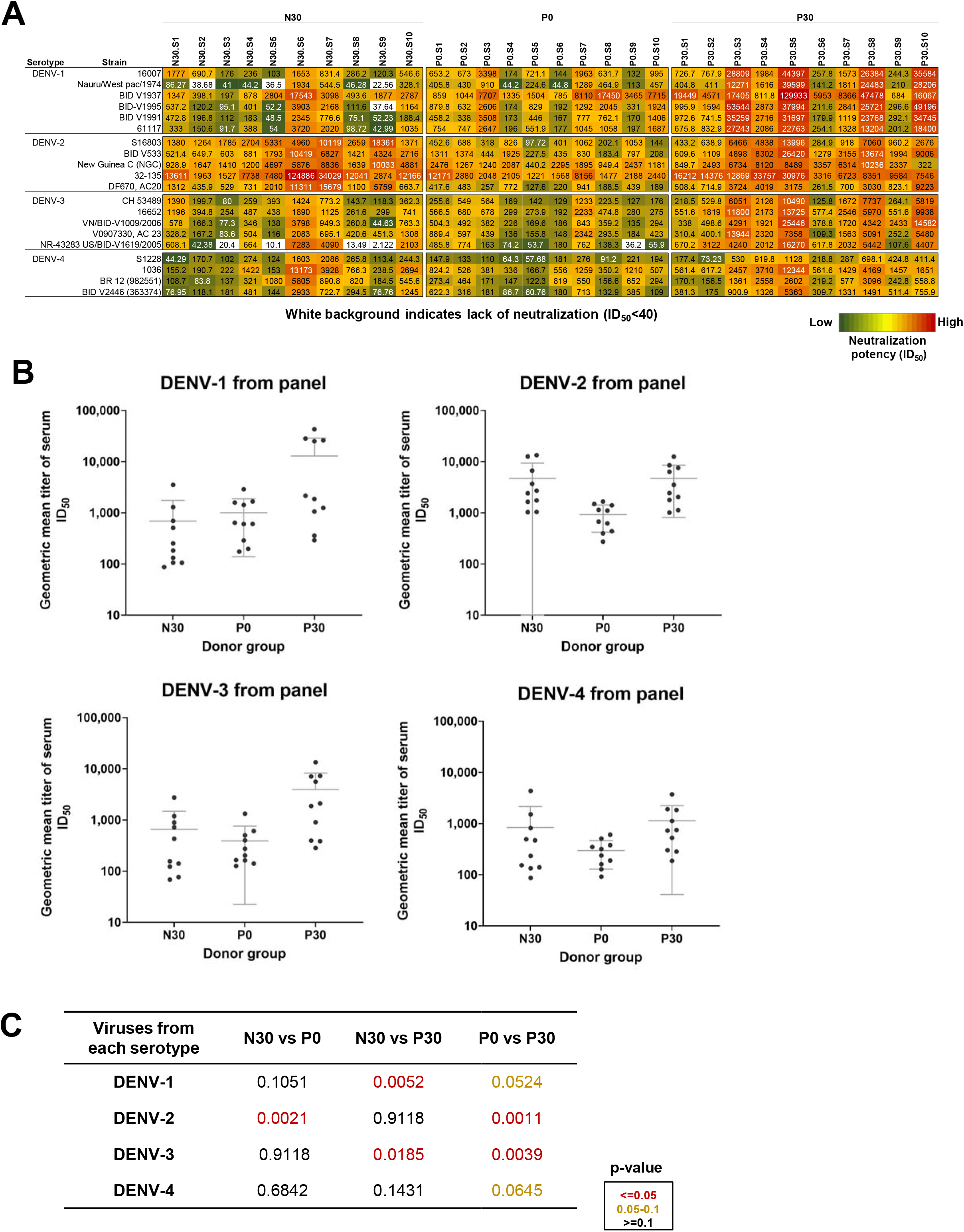
Serum neutralization data. A. Serum-virus neutralization matrix. Shown are the neutralization ID50 values for 30 serum samples against 20 DENV strains. Color scale (in log scale) is green(low)-yellow-red (high) neutralization; ID50 values less than 40 are shown in white. N30: baseline-negative, post-vaccination; P0: baseline-positive, pre-vaccination; P30: P0-matched baseline-positive, post-vaccination. B. Geometric mean titer of serum neutralization ID50. Shown are the log of geometric mean of serum neutralization values of the three sample groups against the four DENV serotypes. Each dot represents the log geometric mean value of a serum/sample against all strains from the respective DENV serotype. C. Shown are the p-values of geometric means ID50 computed between each group for each DENV serotype. p-values ≤0.05 are indicated in red and p-values ≥0.05 and <0.10 are indicated in gold color.

### Effects of TAK-003 vaccination on dengue neutralization by individual donors

To understand the changes in antibody responses between pre- and post-vaccination, the neutralization data for the donor-matched samples from groups P0 and P30 were compared using a variety of strategies. First, for each of the 10 donors, the neutralization potency pre- and post-vaccination was compared for each of the 20 DENVs in the P33700 panel (**Figure 4A**). This analysis revealed that for around half of the donors increased neutralization potency was observed post-vaccination for most or all viruses. For the remaining donors, increased neutralization potency was observed only for a subset of the viruses, with comparable potency observed for most viruses (**Figure 4A**). To further quantify the improvements in neutralization potency in post-vaccination samples, the ID_50_ ratios between post- and pre-vaccination were computed for all strains (**Figure 4B**). A higher ratio would indicate an increase in neutralization potency post-vaccination, compared to pre-vaccination for a given virus. Overall, half of the donors (donors 3, 4, 5, 8, and 10) showed at least a 10-fold improvement in overall neutralization potency post-vaccination. Four additional donors (donors 2, 6, 7, and 9) showed an overall increase in neutralization potency, albeit to a lesser extent and less consistently between strains. Only one donor (donor 1) did not show a substantial change in neutralization potency postvaccination, although substantial per-strain differences in changes in neutralization potency were observed for this donor (**Figure 4B**). For each donor, these results were further categorized by DENV serotype by computing the average of all the post-/pre-vaccination neutralization potency ratios for all strains from the given DENV serotype (**Figure 4C**). Overall, the highest median increase in neutralization potency was observed for DENV serotypes 1 and 3, whereas DENV serotype 4 was associated with the lowest overall increase. Among the 10 donors, the highest increase was observed for donor 5 across all DENV serotypes, whereas the lowest increase was observed for donor 1, with 3 of the 4 DENV serotypes showing an average ratio of less than 1. Together, these results reveal that even though vaccination increased neutralization titer in most donors many of the DENV strains tested in these assays, substantial variability was observed both among donors and among viruses for a given donor.

**Figure 4:**
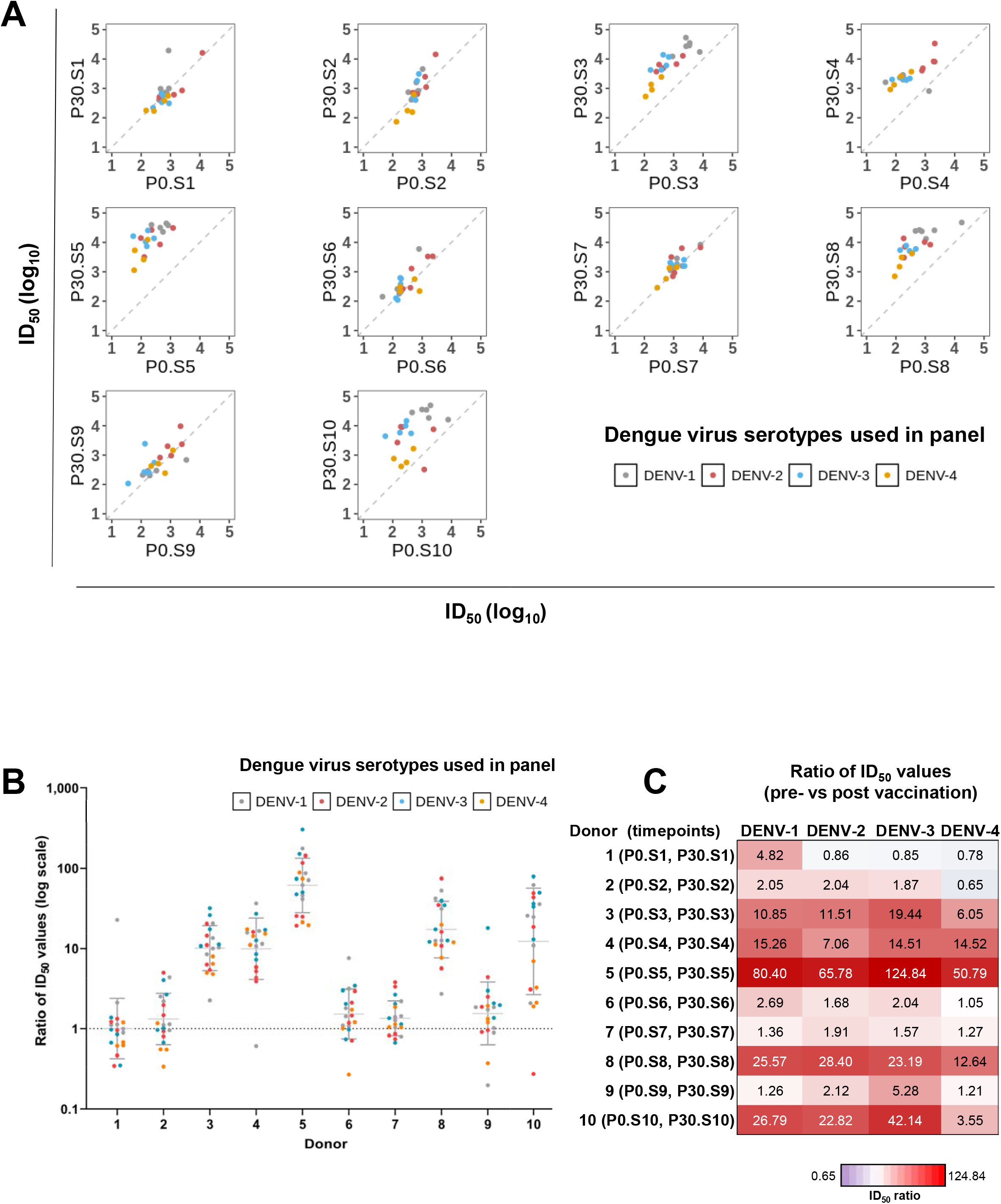
Within-donor changes in neutralizing activity. A. Plot of log ID50 values from pre-(x-axis) vs. post- (y-axis) vaccination. Each plot corresponds to a single donor. Each dot represents the respective log ID50 values for a single strain, and the color represents the respective DENV serotype. B. Ratios of ID50 values between pre- and post-vaccination. For each donor, each dot represents the ID50 ratio value for a single strain, and the color represents the DENV serotype for the respective strain. C. Heatmap of donor neutralization improvement factors by DENV serotype. For each donor and each DENV serotype, shown is the average among the pre/post ID50 ratios for all strains from that DENV serotype. Color scale (log10) is purple (lower ratio)-white-red (higher ratio).

### Impact of baseline seropositivity on vaccine-induced antibody responses

We next sought to understand whether baseline seropositivity resulting from prior DENV virus infection could substantially affect the antibody specificities elicited to subsequent vaccination with TAK-003. To that end, the correlations between the neutralization fingerprints for each pair of serum samples were computed (**Figure 5A**). While high correlations were observed between individual pairs within each of the three sample groups, the correlations within the N30 group appeared to be highest overall, suggesting a convergence of neutralizing antibody specificities in the vaccinated baseline-negative donors (**Figure 5B**). In contrast, the correlations within the P30 group were the lowest, and the correlations between samples from N30 and P0 were overall higher than the correlations between samples from N30 and P30, together suggesting that starting with relatively diverse responses to prior infection in baseline-positive individuals can amplify that diversity post-vaccination (**Figure 5B**). To further interrogate this divergence of antibody responses post-vaccination for the baseline-positive cohort, for each pair of samples, the differences between the correlations from pre- and post-vaccination were computed (**Figure 5C**). Overall, a trend toward lower correlations was observed between donors post-vaccination compared to pre-vaccination, again pointing to an increased divergence of antibody responses post-vaccination, when starting with relatively diverse responses to prior infection.

**Figure 5:**
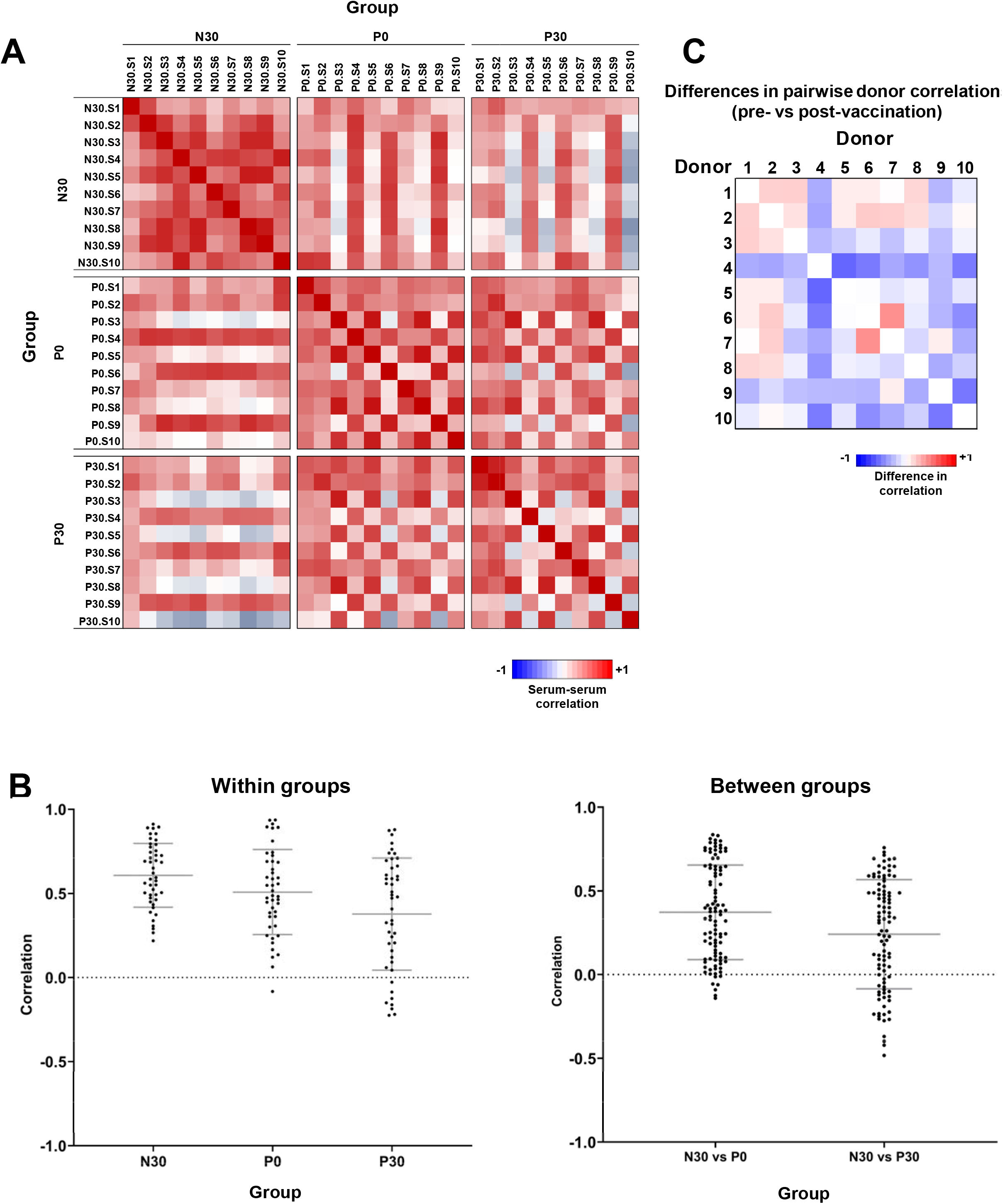
Impact of baseline seropositivity on the specificities of vaccine-induced antibody responses. A. Heatmap of serum-serum neutralization correlation matrix. Shown are correlation coefficients for the neutralization data for each pair of serum samples. Heatmap color scale is blue (−1)-white (0)-red (+1). B. Serum-serum correlation within groups (left) and between groups (right). C. Heatmap of differences in the pairwise donor correlations between pre- (P0) and post- (P30) vaccination. Color scale is blue (lower correlation post-vaccination)-white-red (higher correlation post-vaccination) for each pair of donors.

### NFP deconvolution of polyclonal neutralizing antibody responses to TAK-003 vaccination

To gain insights into the types of antibodies elicited to TAK-003 vaccination, we applied the NFP algorithm to deconvolute the polyclonal neutralization signals (**Figure 3A**) for the three sets of samples (**Figure 6**). For each sample, NFP was applied to predict the overall contribution to polyclonal neutralization by each of the set of 7 reference antibody groups (**Figure 6A**). Different types of antibody specificities were observed for different donors, although samples within each of the three groups N30, P0, and P30 were generally consistent (**Figure 6A**). Overall, in the N30 group, primarily 2D22/3F9-like specificities were predicted to dominate the serum neutralization signals. These results suggest that vaccination in baseline-seronegative individuals resulted in dominant neutralization signal by these DENV-2 specific antibodies, although several subjects also had an NFP signal for cross-reactive (FL-like) antibodies. The dominant NFP signals for samples in the baseline-positive donors (P0 and P30) were predicted to be associated with a diversity of specificities (primarily 1F4/4L5-like DENV-1 and 2D22/3F9-like DENV-2 type-specific; and some FL-like cross-reactive) pre-vaccination, and these signals were generally retained post-vaccination (**Figure 6B**). Overall, NFP signals for 5J7-like, 126/131-like, C10-like, and to some extent 2M2-like specificities were either not observed or infrequently found in any of the groups, suggesting that the epitopes recognized by these types of antibodies may be subdominant compared to other, more potently neutralizing, antibody specificities (**Figure 6A,B**).

**Figure 6:**
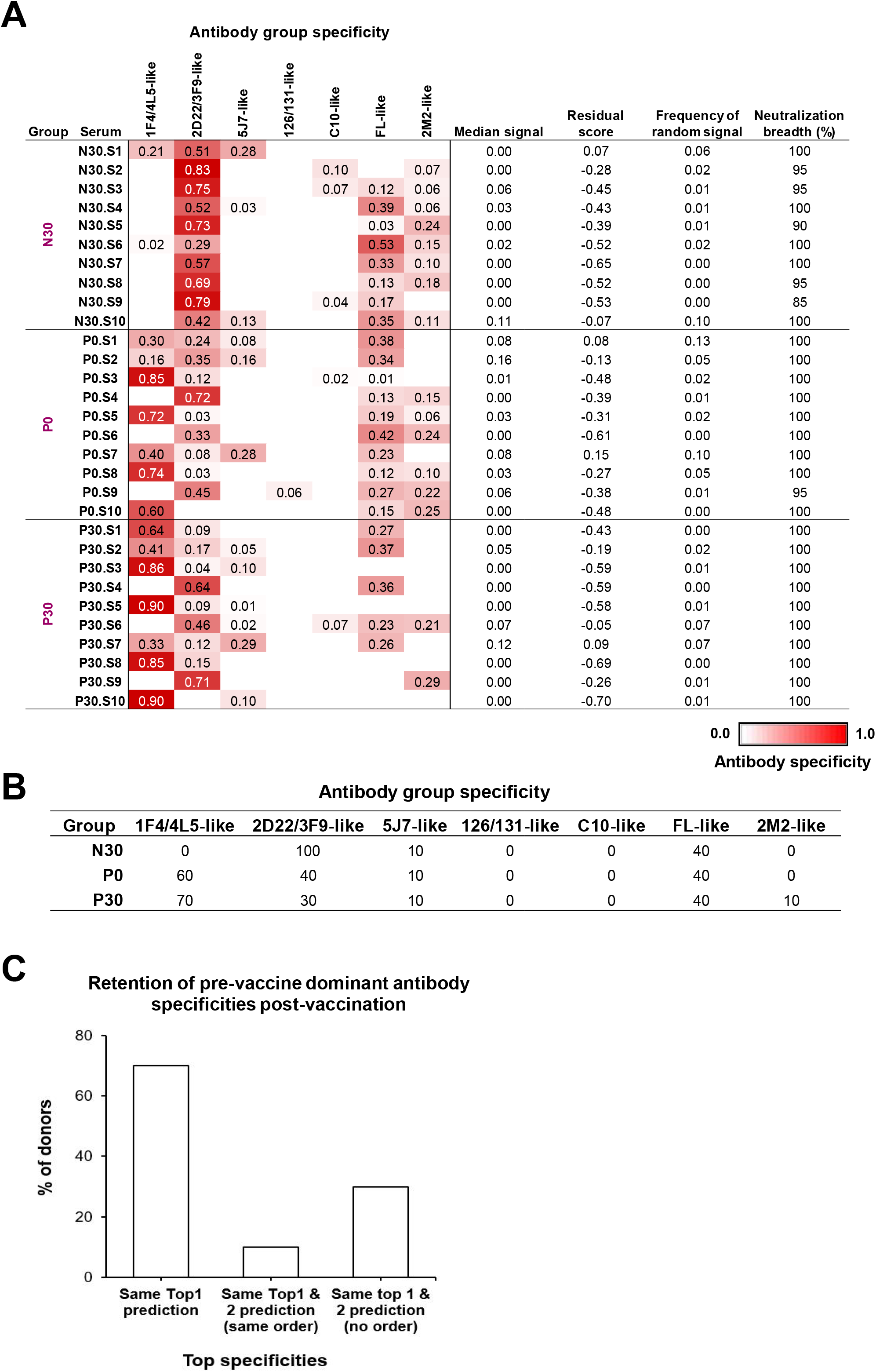
Antibody specificities elicited in response to tetravalent vaccine immunization. A. NFP results for serum samples from all three groups. Column 1 denotes the serum name; columns 2 to 8 are the predicted signals for each of the 7 reference antibody specificities; columns 9 to 11 are related to measures of the reliability of the computational predictions; column 12 shows the neutralization breadth of the given serum sample against the set of 20 strains. B. Sample group-wise antibody specificity frequency. Shown are the frequencies of antibody specificities in each group. C. Retention of pre-vaccine dominant antibody specificities post-vaccination. NFP-predicted specificities were compared between pre- and post-vaccination for each of the 10 donors, and the percent of donors (y-axis) that retained their top specificity (left), and percent of donors that retained both of their top two specificities in the same order (middle) or irrespective of order (right) were computed.

To determine whether there is a general trend related to pre- vs. post-vaccine antibody specificities, the frequency with which donors retained the dominant pre-vaccine specificities after vaccination was analyzed (**Figure 6C**). For 7 of the 10 donors, the same dominant specificity was identified both pre- and post-vaccination, and for 3 of the 10 donors both top two specificities (albeit not necessarily in the same order) were identified both pre- and postvaccination. Together, these results suggest that dominant pre-existing (pre-vaccine) antibody specificities generally were retained post-vaccination.

To further quantify and visualize these findings, the changes in neutralizing antibody specificities pre- vs. post-vaccination within individual donors were analyzed (**Figure 7A**). Overall, although the types of dominant antibody specificities in each donor remained generally unchanged post-vaccination, changes in dominant specificities were observed for certain donors. Further, the differences between the predicted NFP signals pre- and post-vaccination were computed for each donor for each of the reference antibody specificities (**Figure 7B**). Overall, no clear patterns were observed across all donors, with some donors exhibiting an increase in 1F4/4L5-like and/or 2D22/3F9-like specificities, while other donors showed a decrease in 2D22/3F9-like, FL-like, and 2M2-like specificities.

**Figure 7:**
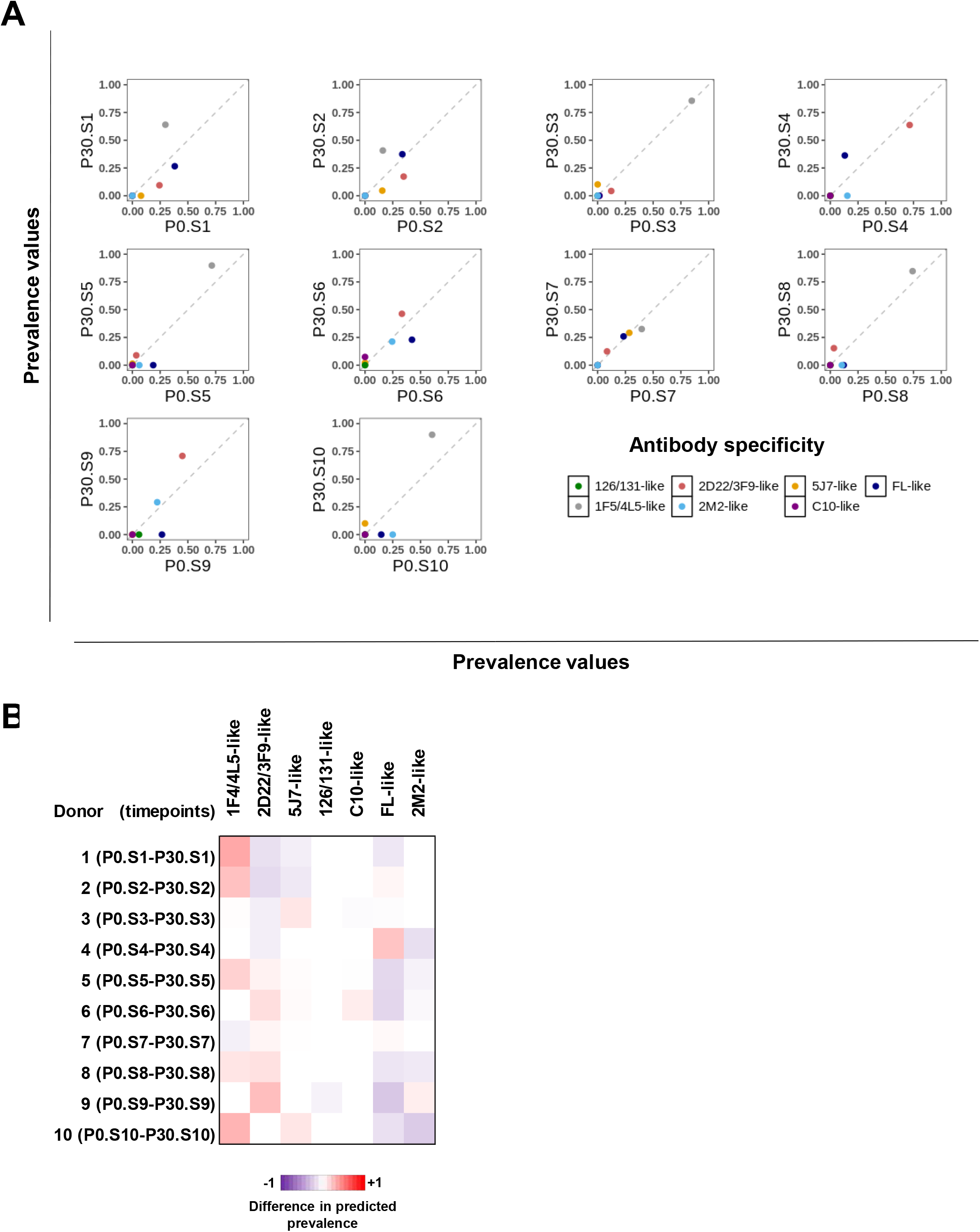
Donor-wise effect of vaccination on antibody specificities. A. Donor-wise comparison of antibody specificities pre- (x-axis) and post- (y-axis) vaccination. Each dot represents the predicted specificity values for each of the reference antibody specificities (colors). B. Donor-wise difference in predicted prevalence of antibody specificities. For each donor (row), shown is a heatmap of the differences for the NFP values for each of the reference antibody specificities (columns). Color scale is blue (−1)-white (0)-red (+1). Red color denotes an increase in the NFP signal for the given antibody specificity post-vaccination, while blue denotes a decrease in signal post-vaccination.

## Discussion

Dengue is one of the major viral diseases lacking an effective vaccine for all populations at risk of disease. A major goal in the dengue virus vaccine field is to develop a vaccine that confers protection against all four DENV serotypes by incorporating representative strains from each serotype. A detailed understanding of antibody responses to these vaccines can help inform further vaccine optimization. Here, we presented the development of a computationally driven approach for deconvoluting polyclonal antibody neutralization into component neutralizing specificities. We applied this approach, called NFP (neutralization fingerprinting), to characterize samples from both baseline-seropositive and baseline-seronegative individuals who had received the tetravalent TAK-003 vaccine, which is currently being evaluated in phase 3 clinical trials.

NFP is a high-throughput method for antibody specificity prediction in samples of interest that had been originally developed in the context of HIV-1 and was adapted here to address the unique characteristics of DENV-specific antibodies, such as serotype specificity. We note that there are certain limitations to this approach. The results and observations from this investigation are based on a limited number of viruses, reference antibodies, and cohort samples. There are more than one serotype-specific epitopes for each DENV serotype, so a larger set of reference antibodies would be required to determine presence or absence of serotype-specific antibodies in a sample. As well, a larger set of reference type-specific antibodies might facilitate use of the NFP method for serological prediction of prior infecting serotype. Further, it is not expected (or even practically possible) to detect more than two epitope specificities in the simulated sera, due to the nature of the algorithm: the sum of all antibody signals for a given serum is 1, and a positive signal is considered >0.25. For the 2-antibody combinations, we could detect both specificities in ~60% of sera, meaning that for a non-negligible proportion of the sera, only the most immunodominant specificity was detected. Observing at least two true positive signals for ~50% of the simulated sera with 4-antibody combinations was lower but generally in line with what we observed for the 2-antibody combinations, suggesting that in the extreme case of adding four dominant antibody specificities, the NFP algorithm could still successfully deconvolute up to two specificities for many of the sera. Related to this, we also point out that while neutralization breadth spanning all four DENV serotypes was observed for virtually all samples (**Figure 3A**), NFP could generally account for the relative contribution to neutralization by the most immunodominant one or two antibody specificities within each individual donor. It is therefore possible that type-specific neutralizing antibodies against, *e.g.*, DENV type 4 exist in a sample, but their signal may be dominated by other more potent specificities, preventing NFP from detecting this signal. In essence, NFP provides an opportunity to understand the most dominant neutralizing antibody specificities in response to infection or multivalent vaccines, such as TAK-003.

The results from this analysis provide a new approach to dissecting immunodominant epitopes of DENV, as well as a potential method to predict prior infecting serotype in polyclonal serum samples from individuals who have experienced DENV infection. Characterizing specificity of neutralizing antibodies following secondary infection, or vaccination of seropositive individuals, has been particularly challenging. This analysis provides new insights into how pre-existing antibodies may shape the evolution of DENV-specific responses to subsequent vaccination or infection (also referred to as immunologic imprinting). A similarity of neutralizing antibody specificities was observed in baseline-negative individuals, suggesting that TAK-003 may elicit generally similar antibody responses in DENV seronegative individuals. In contrast, likely because of the diversity of DENV infections, baseline-seropositive individuals appeared to be less similar in their DENV-specific antibody repertoires pre-vaccination and diverge even more post-vaccination. As no immune correlate of protection against DENV infection has been identified to date, the specificity of neutralizing antibodies capable of protecting against diverse DENVs is not known. DENV-specific antibody repertoires appear to diverge after vaccination of pre-immune individuals, and phase 3 studies demonstrated protective efficacy in this population [11, 19]. In addition, symptomatic disease following 3 or more DENV infections is rare [20, 21]. Thus, it is likely that there is more than one immunological solution to protecting against divergent immune responses.

## Materials and Methods

### Monoclonal antibody neutralization data

Neutralization assays were performed for a set of 19 known DENV-neutralizing antibodies, which included 1F4, 4L5, 2D22, 3F9, 5J7, 126, 131, C10, 1C19, 1M7, 1N5, 1C18, 1L6, 1N8, 3B4, 4E8, 2M2 and 1L12, against a panel of 25 DENV strains, including 7 DENV serotype 1 (DENV-1), 6 DENV-2, 7 DENV-3 and 5 DENV-4 viruses. Neutralization assays for each antibody against a virus strain was repeated 3 times with 8 different concentrations (μg/mL) starting from 100 to 0.00064. Half maximal inhibitory concentration (IC50) values for neutralization of virus in the presence of a given monoclonal antibody were determined by generating neutralization curves using GraphPad software, using virus-only controls to establish maximal virus counts. For the NFP analysis, IC50 values were standardized in the following way. IC50 values of >100 μg/mL were set to 100 as a cutoff for lack of neutralization; selected additional IC50 values that were computed by GraphPad but that corresponded to clear lack of neutralization when the neutralization curves for the respective antibody-virus pair were visualized, were also set to 100. The cutoff of 100 was selected since that was the highest antibody concentration used in the neutralization assays.

In addition, for the neutralization breadth of each antibody, the number of IC50 values less than 100, also was computed. Antibodies neutralizing at least 5 strains were considered for further analysis. A cutoff of 5 strains was used to improve the chances of observing a fingerprinting signal. Specifically, the algorithm relies on analysis of patterns of neutralization; for that to happen, at least a few viruses must be neutralized by each antibody (otherwise, no pattern will exist - all viruses will be resistant). A general cutoff does not exist, but for a virus panel of this small size, we elected to keep antibodies neutralizing at least 5 viruses. With larger virus panel sizes, we will increase that cutoff, since having too few neutralized viruses per antibody can lead to insufficient data for the algorithm to analyze, and to less reliable predictions. Among the 19 antibodies, antibody 5C8, which did not neutralize any strain among the 25, was excluded from the dataset and further analysis. The other 18 antibodies, with neutralization breadth of 20 to 100%, were selected for further analysis and the resulting antibody-virus neutralization matrix is shown in Figure 1A.

### Generation of different size of viral panel candidates

In our previous studies on HIV-1, we showed the selection of an optimized panel can lead to significantly improved prediction accuracy [17]. Since the selection of strains and panel size are important for NFP prediction accuracy, we searched for candidate virus panels of size 18 to 22. All possible combinations of virus strains for each panel size were generated and the panels with less than 4 strains from any of the DENV types were excluded from further consideration. In other words, only panels including at least 4 strains from each dengue type were included in the subsequent analysis. The final number of panels included 149,450 18-strain panels, 86,590 19-strain panels, 34,916 20-strain panels, 10,080 21-strain panels and 2,070 22-strain panels. In addition, we also evaluated the prediction accuracy of the full 25-strain panel, as a comparison and benchmark for improvement.

### Computational simulation of polyclonal serum neutralization with two-antibody combinations

To select an optimized panel from the list of generated viral panels, we need to be able to compare the prediction accuracy of different size of virus panel candidates on serum samples. As the real human samples are not suitable for use as a benchmark because of the unknown composition and prevalence of antibody specificities, we developed a framework for generating simulated polyclonal serum neutralization data, with pre-defined composition and prevalence of antibody specificities. To identify optimized virus panels, a set of simulated sera were generated with the combinations of two antibody specificities. Sera data were simulated using two-antibody combinations of neutralization data that mixed an equal amount of an antibody representative from any two of the 7 antibody groups, for all 25 strains, and allowing for an antibody potency scaling factor and experimental noise, for the final serum neutralization computation. For two-antibody combinations using 7 different groups with 21 pairwise combinations, 2,100 simulated sera (100 sera/combination) were generated. The simulated serum neutralization data is used as a quantitative benchmark for the comparison and evaluation of the prediction ability of the NFP algorithm for different candidate virus panels.

### Computational simulation of polyclonal serum neutralization with four dengue type-specific antibody combinations

One of the challenges for DENV analysis, compared to HIV-1, is the existence of antibody type-specificity. Therefore, an additional important question is whether the selected panel with the fingerprinting algorithm will be able to distinguish neutralization breadth that is due to polyclonal breadth resulting from a combination of type-specific monoclonal antibodies from monoclonal neutralization breadth. To address this question, we generated simulated sera by combining one antibody from each of the four type-specific antibody groups. The goal of these experiments is to determine whether the algorithm can successfully predict that there are mixtures of type-specific antibodies, rather than erroneously indicate the presence of broad monoclonal antibodies. This is an extreme case scenario (with mixtures of four different specificities), and it is not expected (or even possible) to be able to correctly identify the presences of all four DENV serotype-specific signals at a time. Rather, we aim to determine the levels of false positives for the different cross-neutralizing antibody groups. To that end, simulated serum data were generated using combinations of four type-specific antibodies that mixed equal amount of an antibody representative for 4 DENV-specific groups. During the simulation, if one or more antibodies do not neutralize a strain (IC50 value ≥ 100), then the neutralization values of the other antibodies are considered for the simulation. A total of 1,000 simulated serum data was generated and used as a benchmark dataset to select an optimized virus panel.

### Serum neutralization data

Neutralization experiments were performed for 30 serum samples from 20 donors. Samples N30.S1 to N30.S10 were collected 30 days after vaccination from 10 subjects with no history of previous DENV infection (baseline negative; N). A second set of 10 subjects with prior DENV infection (baseline positive; P) was included, and the samples were collected at 2 time points. Samples P0.S1 to P0.S10 were collected at day 0 before vaccination (pre-vaccination; P0) and samples P30.S1 to P30.S10 were collected at 30 days after vaccination (post-vaccination, P30). Serum neutralization experiments were conducted against the selected viral panel and the assay for each serum against a virus strain was repeated 3 times with 8 different dilutions starting from 1/10 to 1/163840. ID50 values for neutralization of a given virus strain in the presence of serum sample were determined by generating neutralization curves using GraphPad software, using virus-only controls to establish maximal virus counts. The resulting serum-virus neutralization matrix is shown in Figure 3A.

---

Supplementary figure 1:

A. Large-scale simulation of polyclonal antibody neutralization with two antibody specificities. The first panel heatmap represents combinations of two antibody specificities from the 7 dengue antibody groups. The middle panel represents the resulting simulated serum neutralization data, shown as a potency-based heatmap (red-yellow-green for most to least potent). The third panel represents the number of possible combinations and final number of simulated sera data.

B. Large-scale simulation of polyclonal antibody neutralization with four antibody specificities. The first panel heatmap represents combinations of four antibody specificities from the four DENV serotype-specific antibody groups. The middle panel represents the resulting simulated serum neutralization data. The third panel represents the number of possible combinations and final number of simulated sera data.

C. Bar graphs of true positives (red) and false positives (blue) for the 25-strain (left) and selected 20-strain (right) panels against simulated sera with 2-antibody combinations.

## References

1. World Health Organization, Dengue vaccine: WHO position paper, July 2016 - recommendations. Vaccine, 2017. 35(9): p. 1200–1201.

2. Bhatt, S., et al., The global distribution and burden of dengue. Nature, 2013. 496(7446): p. 504–507.

3. Messina, J.P., et al., Global spread of dengue virus types: mapping the 70 year history. Trends in microbiology, 2014. 22(3): p. 138–146.

4. Halstead, S.B., Dengue Antibody-Dependent Enhancement: Knowns and Unknowns. Microbiol Spectr, 2014. 2(6).

5. Katzelnick, L.C., et al., Antibody-dependent enhancement of severe dengue disease in humans. Science, 2017. 358(6365): p. 929–932.

6. Clements, D.E., et al., Development of a recombinant tetravalent dengue virus vaccine: immunogenicity and efficacy studies in mice and monkeys. Vaccine, 2010. 28(15): p. 2705–15.

7. George, S.L., et al., Safety and Immunogenicity of a Live Attenuated Tetravalent Dengue Vaccine Candidate in Flavivirus-Naive Adults: A Randomized, Double-Blinded Phase 1 Clinical Trial. J Infect Dis, 2015. 212(7): p. 1032–41.

8. Capeding, M.R., et al., Clinical efficacy and safety of a novel tetravalent dengue vaccine in healthy children in Asia: a phase 3, randomised, observer-masked, placebo-controlled trial. Lancet, 2014. 384(9951): p. 1358–65.

9. Villar, L., et al., Efficacy of a Tetravalent Dengue Vaccine in Children in Latin America. New England Journal of Medicine, 2014. 372(2): p. 113–123.

10. Hadinegoro, S.R., et al., Efficacy and Long-Term Safety of a Dengue Vaccine in Regions of Endemic Disease. New England Journal of Medicine, 2015. 373(13): p. 1195–1206.

11. Biswal, S., et al., Efficacy of a Tetravalent Dengue Vaccine in Healthy Children and Adolescents. New England Journal of Medicine, 2019. 381(21): p. 2009–2019.

12. Rivera, L., et al., Three years efficacy and safety of Takeda’s dengue vaccine candidate (TAK-003). Clin Infect Dis, 2021.

13. Michlmayr, D., et al., Characterization of the Type-Specific and Cross-Reactive B-Cell Responses Elicited by a Live-Attenuated Tetravalent Dengue Vaccine. J Infect Dis, 2021. 223(2): p. 247–257.

14. Swanstrom, J.A., et al., Analyzing the Human Serum Antibody Responses to a Live Attenuated Tetravalent Dengue Vaccine Candidate. J Infect Dis, 2018. 217(12): p. 1932–1941.

15. Georgiev, I.S., et al., Delineating antibody recognition in polyclonal sera from patterns of HIV-1 isolate neutralization. Science, 2013. 340(6133): p. 751–6.

16. Raju, N., I. Setliff, and I.S. Georgiev, NFPws: a web server for delineating broadly neutralizing antibody specificities from serum HIV-1 neutralization data. Bioinformatics, 2019. 35(18): p. 3502–3504.

17. Doria-Rose, N.A., et al., Mapping Polyclonal HIV-1 Antibody Responses via Next-Generation Neutralization Fingerprinting. PLoS Pathog, 2017. 13(1): p. e1006148.

18. Gallichotte, E.N., et al., Human dengue virus serotype 2 neutralizing antibodies target two distinct quaternary epitopes. PLoS Pathog, 2018. 14(2): p. e1006934.

19. Biswal, S., et al., Efficacy of a tetravalent dengue vaccine in healthy children aged 4-16 years: a randomised, placebo-controlled, phase 3 trial. Lancet, 2020. 395(10234): p. 1423–1433.

20. Chinnawirotpisan, P., et al., Detection of concurrent infection with multiple dengue virus serotypes in Thai children by ELISA and nested RT-PCR assay. Arch Virol, 2008. 153(12): p. 2225–32.

21. Castellanos, J.E., et al., Description of high rates of unapparent and simultaneous multiple dengue virus infection in a Colombian jungle settlement. Trop Biomed, 2016. 33(2): p. 375–382.

