## Supplemental Figure 1 for "Neutralization Fingerprinting Technology for Characterizing Polyclonal Antibody Responses to Dengue Vaccines"

### Slide 1
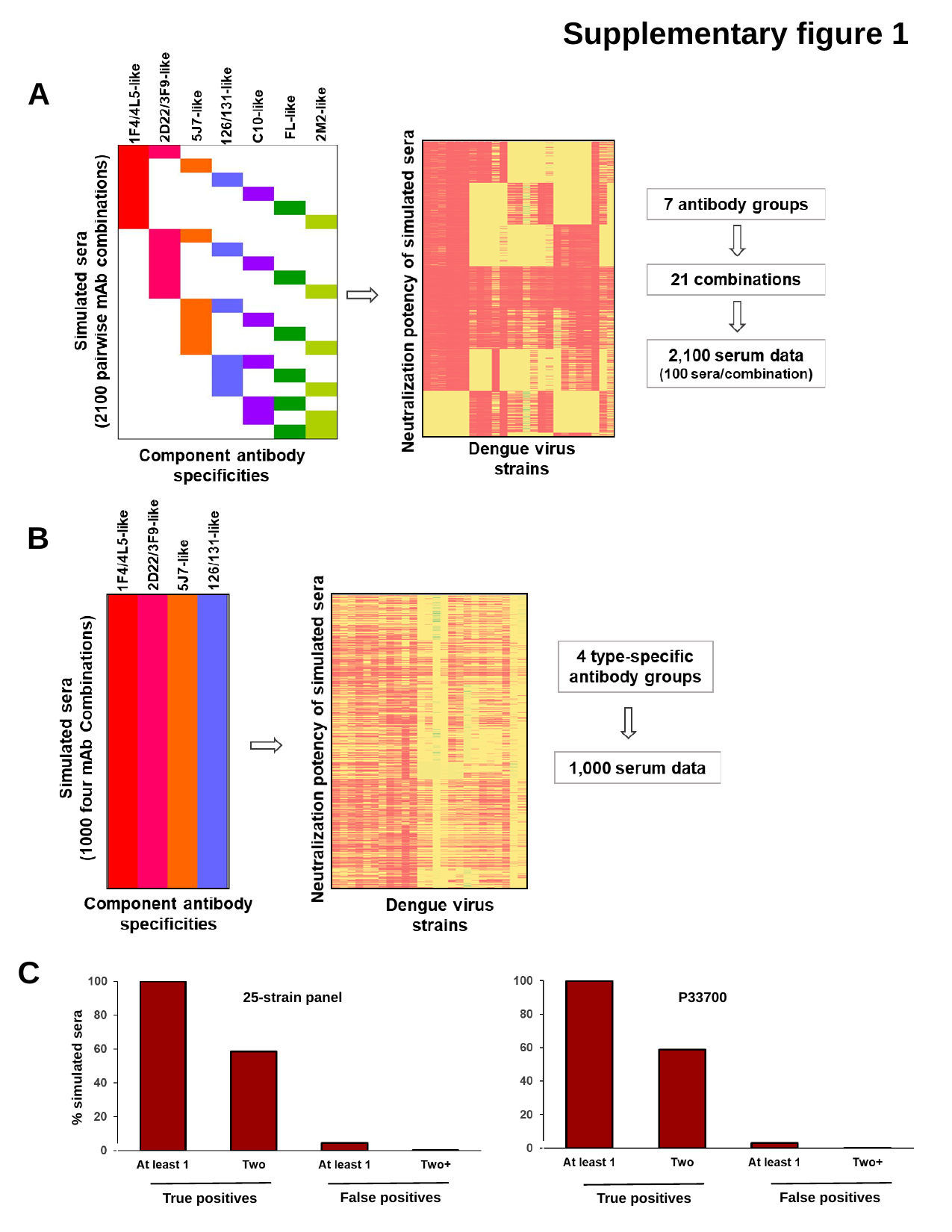

Supplementary figure 1
A
B
C
25-strain panel
P33700
% simulated sera
False positives
False positives
True positives
True positives
